# Chromatin remodelers couple inchworm motion with twist-defect formation to slide nucleosomal DNA

**DOI:** 10.1101/297762

**Authors:** Giovanni B. Brandani, Shoji Takada

## Abstract

ATP-dependent chromatin remodelers are molecular machines that control genome organization by repositioning, ejecting, or editing nucleosomes, activities that confer them essential regulatory roles on gene expression and DNA replication. Here, we investigate the molecular mechanism of active nucleosome sliding by means of molecular dynamics simulations of the Snf2 remodeler in complex with a nucleosome. During its inchworm motion driven by ATP consumption, the remodeler overwrites the original nucleosome energy landscape via steric and electrostatic interactions to induce sliding of nucleosomal DNA unidirectionally. The sliding is initiated at the remodeler binding location via the generation of twist defects, which then spontaneously propagate to complete sliding throughout the entire nucleosome. We also reveal how remodeler mutations and DNA sequence control active nucleosome repositioning, explaining several past experimental observations. These results offer a detailed mechanistic picture of remodeling important for the complete understanding of these important biological processes.

## INTRODUCTION

Eukaryotic genomes are compacted into the cell nucleus via the formation of nucleosomes, each of them consisting of ~147 base pairs (bp) of DNA wrapping around a protein histone octamer(Richmond & Davey, 2003). After having been initially considered as passive building blocks of chromatin organization, nucleosomes became to be recognized as active regulators of DNA transcription and replication(Lai & Pugh, 2017). An origin of this regulation is the steric effect that inhibits other DNA-binding proteins, such as transcription factors, from accessing nucleosomal DNA(Field et al., 2008), suggesting the requirement of fine control of nucleosome positioning along the genomic sequence(Lai & Pugh, 2017; Struhl & Segal, 2013). For instance, repositioning of nucleosomes located next to transcription start sites enables the dynamic regulation of gene expression in response to stresses such as heat shock(Reja, Vinayachandran, Ghosh, & Pugh, 2015; Shivaswamy et al., 2008). While *in vitro* the nucleosome locations are solely determined by DNA mechanics(Freeman, Lequieu, Hinckley, Whitmer, & de Pablo, 2014; Morozov et al., 2009), precise positioning *in vivo* is largely controlled by chromatin remodelers(Lai & Pugh, 2017; Struhl & Segal, 2013), which are ATP-dependent molecular machines(Clapier, Iwasa, Cairns, & Peterson, 2017; Zhou, Johnson, Gamarra, & Narlikar, 2016). High-resolution structures of some of these remodelers bound to nucleosomes were obtained very recently by cryo-EM(Ayala et al., 2018; Eustermann et al., 2018; Farnung, Vos, Wigge, & Cramer, 2017; Liu, Li, Xia, Li, & Chen, 2017). While these static structures provide crucial insights, currently missing are the dynamic aspects of how these molecular machines work, on which the current study focus by molecular dynamics simulations.

Remodelers are molecular motors that consume ATP to perform a wide variety of functions related to genome organization(Clapier et al., 2017; Zhou et al., 2016): facilitating nucleosome assembly(Torigoe et al., 2011) and precise spacing(Gkikopoulos et al., 2011; Krietenstein et al., 2016; Yang, Madrid, Sevastopoulos, & Narlikar, 2006), controlling DNA accessibility via nucleosome sliding(Harada et al., 2016) or histone ejection(Boeger, Griesenbeck, Strattan, & Kornberg, 2004), and nucleosome editing via exchange between different histone variants(Bruno et al., 2003). These activities enable remodelers to maintain chromatin organization after disruptive events such as replication(Lai & Pugh, 2017; Vasseur et al., 2016), and to regulate gene expression via the dynamic control of nucleosome positions(Boeger et al., 2004; Lai & Pugh, 2017; Lorch & Kornberg, 2015; Reja et al., 2015; Shivaswamy et al., 2008).

Although the changes in chromatin organization induced by remodelers have been widely documented(Gkikopoulos et al., 2011; Krietenstein et al., 2016; Lai & Pugh, 2017), the precise molecular mechanisms are still far from being clear(Clapier et al., 2017; Mueller-Planitz, Klinker, & Becker, 2013; Zhou et al., 2016). The complexity comes in part from the existence of a wide variety of remodelers with different structures and functions(Zhou et al., 2016), which has led to several possible classifications into remodeler sub-families(Flaus & Owen-Hughes, 2011). Each remodeler consists of many distinct domains, which act in concert to confer specificity to the remodeling activity (e.g. nucleosome sliding vs ejection)(Clapier et al., 2017; Zhou et al., 2016) and to fine-tune it via substrate recognition (e.g. of histone tail modifications)(R. Blossey & Schiessel, 2008; Narlikar, 2010). Despite this complexity, all remodelers share a conserved translocase domain(Clapier et al., 2017): an ATPase motor capable of unidirectional sliding along DNA via binding and hydrolysis of ATP between its two RecA-like lobes, structurally similar to those found in helicases(Flaus & Owen-Hughes, 2011; Zhou et al., 2016). The translocase domain of most remodelers binds nucleosomes at the superhelical location (SHL) 2(Clapier et al., 2017; Farnung et al., 2017; Liu et al., 2017), i.e. two DNA turns away from the dyad symmetry axis (SHL 0)(Luger, Mäder, Richmond, Sargent, & Richmond, 1997) (Fig. 1a). Many remodelers induce sliding of nucleosomal DNA towards the dyad from the translocase binding location(Brahma et al., 2017; Saha, Wittmeyer, & Cairns, 2005; Schwanbeck, Xiao, & Wu, 2004; Zofall, Persinger, Kassabov, & Bartholomew, 2006). This may represent a shared fundamental mechanism at the basis of most remodeling activities; the interactions with the additional domains would then confer specificity to the remodeler, allowing for substrate recognition and determining whether the final outcome is nucleosome repositioning, histone ejection or histone exchange(Clapier et al., 2017). For instance, recent experiments suggested that the INO80 remodeler causes histone exchange by sliding nucleosomal DNA from its translocase binding site around SHL 6(Ayala et al., 2018; Brahma et al., 2017; Eustermann et al., 2018). Therefore, the detailed characterization of active nucleosome sliding by the translocase domain would represent a significant step forward in our understanding of chromatin remodeling.

There is much experimental evidence suggesting that the translocase domain of remodelers, as well as some helicases, slides unidirectionally along DNA via an inchworm mechanism(Clapier et al., 2017; Velankar, Soultanas, Dillingham, Subramanya, & Wigley, 1999), which may also be viewed as a molecular ratchet(Farnung et al., 2017; Gu & Rice, 2010), processing by 1 bp every ATP cycle. A minimal inchworm model requires 3 distinct chemical states, apo, ATP-bound, and ADP-bound, which are coupled to conformational changes of the translocase (Fig. 1b). The two lobes are either distant in the open form or in contact in the closed form. On top, conformational changes modulate interaction strengths with DNA(Gu & Rice, 2010). To translocate along DNA in the direction from lobe 1 to lobe 2 (which would correspond to sliding nucleosomal DNA in the direction from SHL 2 towards the dyad, as in most remodelers(Liu et al., 2017)), ATP binding to the translocase first induces the transition from an open to a closed conformation (first and second cartoons in Fig. 1b). During closure, lobe 2-DNA interactions are stronger than those of lobe 1, so that lobe 1 will detach from the DNA and move towards lobe 2 by 1 bp, which maintains its position. Then, ATP hydrolysis is accompanied by weakening of the lobe 2 interactions with DNA relative to those of lobe 1, so that during the conformational change from the closed to the open state lobe 2 now moves away from lobe 1 by 1 bp (third cartoon in Fig. 1b). The cycle is then completed by the release of ADP and the change of the interaction strengths to their initial apo-state values. This mechanism was firstly suggested for helicases from their crystal structures that show ATPase closure and changes in lobes-DNA interactions as a function of the chemical state(Gu & Rice, 2010; Velankar et al., 1999). For remodelers, the same mechanism was suggested based on the structural similarity to helicases(Flaus & Owen-Hughes, 2011; Liu et al., 2017; Zhou et al., 2016) and the recent cryo-EM structures of nucleosome-bound remodelers in the open and closed conformations(Farnung et al., 2017; Liu et al., 2017). However, while the inchworm model can explain the motion of remodelers along naked DNA(Sirinakis et al., 2011), its application to nucleosome repositioning is far from trivial, since such motion would eventually result into steric clashes between the remodeler and the histone octamer. Furthermore, complete nucleosome repositioning necessarily involves breakage of the many histone-DNA contacts that stabilize the nucleosome structure(Luger et al., 1997), and it is not clear how the remodeler may perturb the contacts far away from the binding location at SHL 2.

**Figure 1:**
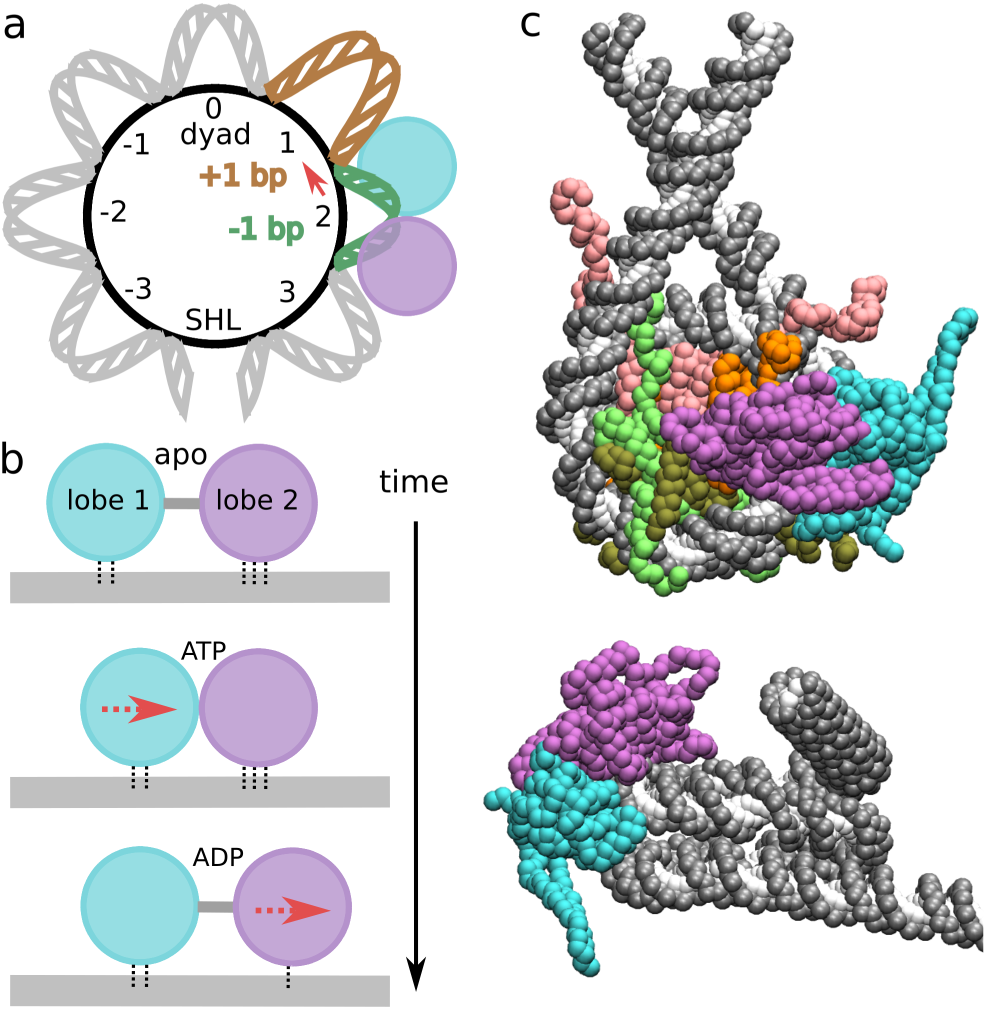
Structure of the translocase-nucleosome complex. (a) Cartoon where nucleosome regions are indicated by the number of DNA turns from the dyad symmetry axis; superhelical location (SHL) 0 corresponds to the dyad, the ATPase domain (lobe 1 in cyan, lobe 2 in purple) binds DNA at SHL 2, the strong histone-DNA contact points are located at the half-integer SHLs where the DNA minor groove faces the octamer (for clarity, we only depict the region from SHL −4 to +4). To analyze DNA sliding, we track the base pair indexes Δbp_i_ at these contact points relative to the initial conformation. If, for example, nucleosome sliding starts with the motion of DNA at contact point 1.5 by 1 bp towards the dyad (red arrow), the contact point Δbp_1.5_ will increase from ~0 to ~1, and this will also indicate the formation of an over-twist defect at SHL 1 (brown) and an undertwist defect at SHL 2 (green), which now accommodate respectively an extra and a missing base pair relative to the reference nucleosome conformation in the crystal structure with PDB id 1KX5. (b) Inchworm mechanism of translocase motion along DNA. ATP binding induces a conformational change from open to closed, with the motion of lobe 1 towards lobe 2 by 1 bp, due to the weaker DNA contacts of the former. ATP hydrolysis weakens the lobe 2-DNA contacts and induces opening via the motion of lobe 2 away from lobe 1 by 1 bp. ADP release completes the cycle. (c) Two views of the initial Snf2-nucleosome structure for our coarse-grained MD simulations: DNA backbone (phosphate and sugar groups) in gray, bases in white, translocase lobes in cyan and purple, histones H3 in pink, H4 in orange, H2A and H2B in light and dark green respectively (in top snapshot only).

Experimental studies have also highlighted many diverse structural changes of the nucleosome occurring during remodeling, such as DNA twisting(Mueller-Planitz et al., 2013), loops(Y. Zhang et al., 2006) or histone deformations(Sinha, Gross, & Narlikar, 2017), suggesting these may be directly responsible for nucleosome sliding. Interestingly, similar structural changes are also believed to mediate spontaneous sliding(Ralf Blossey & Schiessel, 2011). Therefore, insights from research on spontaneous nucleosome repositioning may shed light into the more complex active case. Indeed, DNA sliding on nucleosomes can also be simply driven by thermal fluctuations(Meersseman, Pennings, & Bradbury, 1992). Modes of nucleosome repositioning can be classified in two types depending on whether sliding is accompanied by the rotation of DNA around its axis(Niina, Brandani, Tan, & Takada, 2017). In the rotation-uncoupled mode, sliding proceeds via large steps of about a DNA turn (~10 bp)(Lequieu, Schwartz, & de Pablo, 2017; Niina et al., 2017; Schiessel, Widom, Bruinsma, & Gelbart, 2001), possibly facilitated by the formation of loops(Lequieu et al., 2017; Schiessel et al., 2001). On the other hand, in the rotation-coupled mode, DNA sliding proceeds at small steps of 1 bp via a screw-like motion(Niina et al., 2017), facilitated by the formation of twist defects(Brandani, Niina, Tan, & Takada, 2018; Edayathumangalam, Weyermann, Dervan, Gottesfeld, & Luger, 2005; Kulić & Schiessel, 2003; Richmond & Davey, 2003; Shaytan et al., 2016; van Holde & Yager, 1985). Twist defects are the structural deformations of DNA allowing to accommodate different numbers of base pairs between the strong histone-DNA contact points, which correspond to half-integer SHL locations(Edayathumangalam et al., 2005; Luger et al., 1997). Between two adjacent contact points, while canonical DNA turns contains ~10-bp, 9 bp (a missing bp) or 11 bp (an extra bp) may also be accommodated, corresponding to deformations referred to as under-twist and over-twist defects respectively (see an example illustrated in Fig. 1a, where the anti-clockwise motion of DNA at the contact point at SHL 1.5 (the red arrow) generates DNA under-twisting at SHL 2 (green) and DNA over-twisting at SHL 1 (brown)). The spontaneous formation and propagation of twist defects around the nucleosome causes repositioning by 1 bp at the time(Brandani et al., 2018; Kulić & Schiessel, 2003). Notably, the small characteristic step sizes observed during nucleosome sliding by the ISWI and RSC remodelers(Deindl et al., 2013; Harada et al., 2016) suggests the role of DNA twisting in the molecular mechanism(Mueller-Planitz et al., 2013). Furthermore, having the same step size, active sliding via twist defects is compatible with the inchworm motion of the translocase domain. While some remodelers have been shown to induce the formation of large DNA loops in nucleosomes(Y. Zhang et al., 2006), this may be due to interactions with extra domains in the remodeler, and their presence should not rule out the importance of twist defects in chromatin remodeling.

Due to the ubiquitous importance of remodelers for chromatin organization, gene expression and replication(Lai & Pugh, 2017), the detailed understanding of their molecular mechanism of action would be extremely valuable. Coarse-grained molecular dynamics (MD) simulations(Takada et al., 2015) represent an ideal tool for approaching this problem, since their resolution can be high enough to accurately represent the potential key steps occurring during active nucleosome repositioning(Brandani et al., 2018; Flechsig & Mikhailov, 2010; Niina et al., 2017), while achieving a speed-up of several orders of magnitude relative to all-atom simulations(Takada et al., 2015), for which the system size and the relevant time scales would result in an exceedingly large computational cost. Notably, coarse-graining approaches have been successfully applied to the study of nucleosome dynamics(Chang & Takada, 2016; Freeman, Lequieu, et al., 2014; Kenzaki & Takada, 2015; B. Zhang, Zheng, Papoian, & Wolynes, 2016), including spontaneous repositioning(Brandani et al., 2018; Lequieu et al., 2017; Niina et al., 2017), and ATP-dependent molecular motors(Flechsig & Mikhailov, 2010; Koga & Takada, 2006).

In this work, we investigate the fundamental mechanism of active repositioning in a minimal system consisting of the ATPase-translocase domain of the Snf2 remodeler from yeast in complex with the nucleosome(Liu et al., 2017). Firstly, we test that our model can reproduce the expected inchworm mechanism and unidirectional sliding during ATP consumption. By comparing to the spontaneous case, we show how the remodeler induces directed repositioning by modifying the nucleosome free energy landscape via steric effects and long-range electrostatic interactions, explaining past experimental data on Snf2 mutants(Liu et al., 2017). Nucleosome repositioning occurs by coupling the ATPase inchworm motion to the formation and propagation of twist defects starting from the remodeler binding location at SHL 2. Finally, we reveal how DNA sequence can be exploited to control the kinetics of the system, consistently with its role in determining the repositioning outcome of many remodelers(Krietenstein et al., 2016; Lorch, Maier-Davis, & Kornberg, 2014; Winger & Bowman, 2017).

## RESULTS

### MD simulations of remodeler translocase sliding on naked and nucleosomal DNA via an inchworm mechanism

We performed coarse-grained MD simulations of the ATP-dependent translocase domain of the Snf2 remodeler both on naked DNA and when bound to nucleosomes (Fig. 1c). The nucleosome model is the same as that previously employed to study spontaneous nucleosome repositioning(Brandani et al., 2018; Niina et al., 2017), whereas the remodeler model and its interactions with the DNA are based on the cryo-EM structure of the Snf2-nucleosome complex(Liu et al., 2017). Our computational model coarse-grains proteins at the level of individual residues(Li, Wang, & Takada, 2014) and DNA at the level of sugar, phosphate, and base groups, capturing the sequence-dependent flexibilities of base steps(Freeman, Hinckley, Lequieu, Whitmer, & de Pablo, 2014; Olson, Gorin, Lu, Hock, & Zhurkin, 1998) (see the Materials and Methods section and Refs. (Freeman, Hinckley, et al., 2014; Li et al., 2014; Niina et al., 2017) for more details). In most simulations, we used a 2-bp periodic sequence formed by repeating ApG base steps (polyApG). This sequence was chosen because it was shown to display an intermediate flexibility on the nucleosome(Brandani et al., 2018). We also investigated the effect on active repositioning due to strong nucleosome positioning sequences(Lowary & Widom, 1998; Segal et al., 2006), and the introduction of sequence motifs with different flexibilities at target regions.

Based on the inchworm model (Fig. 1b), each remodeler chemical state (apo, ATP or ADP) corresponds to slightly different force-field parameters of the coarse-grained potential, and we simulate an ATP cycle (apo→ATP→ADP→apo) via switching the potential during the MD simulation, a common strategy in coarse-grained studies of molecular motors(Koga & Takada, 2006; Yao, Kenzaki, Murakami, & Takada, 2010). Initially, in the open apo state, the remodeler configuration and the strengths of the interactions between ATPase lobes and DNA are as found in the cryo-EM structure with PDB id 5X0Y(Liu et al., 2017); then, switching to the ATP-bound potential enhances the attraction between the two lobes, favoring the closed conformation of the remodeler. ATP hydrolysis is emulated by switching to the ADP-bound potential, which reduces the lobe 2-DNA interactions by a factor of 0.8 and weakens the attractive interaction between the two lobes to favor the open conformation. In all our MD trajectories (40 on naked DNA, 100 on nucleosome), switching from apo to ATP states occurs at time 0 after 2x10^7^ MD equilibration steps in the open conformation, ATP hydrolysis occurs after 10^7^ MD steps, which are sufficient for the full relaxation of the system in the closed conformation, and finally switching back to the apo state occurs after 10^7^ steps (which are sufficient to observe translocase opening). More details on the simulation protocol are provided in the Materials and Methods section.

To analyze the repositioning dynamics in our MD trajectories, we track the motion of the DNA base pairs relative to the two individual ATPase lobes and relative to the histone octamer at the 14 histone-DNA contact points, located at the half-integer SHLs where the DNA minor groove faces the octamer. We refer to these collective variables as the contact indexes: Δbp_L1_ and Δbp_L2_ for the remodeler lobes and Δbp_i_ for the histone contacts, where i is the half-integer valued SHL of the contact (these contacts will be indicated by their SHL value, e.g. contact point 1.5). As in our previous work(Brandani et al., 2018), the nucleosome contact indexes are evaluated relative to the 147-bp conformation found in the crystal structure with PDB id 1KX5(Davey, Sargent, Luger, Maeder, & Richmond, 2002), which does not display twist defects. Using these contact indexes, we can fully characterize the remodeler’s inchworm dynamics, the sliding of DNA in the nucleosome, and the potential role played by twist defects. These DNA deformations are distributed around integer-valued SHLs lacking direct histone-DNA contacts, and can either involve DNA over-twisting accommodating an extra base pair relative to the 1KX5 reference (over-twist defect), or DNA under-twisting accommodating a missing base pair (under-twist defect)(Brandani et al., 2018; Kulić & Schiessel, 2003; Richmond & Davey, 2003; Richmond & Widom, 2000; van Holde & Yager, 1985). A twist defect at SHL i can be simply evaluated by the difference between the neighboring contact indexes (i-1/2 and i+1/2): a defect value close to zero corresponds to the standard non-defect case, a value close to 1 to an over-twist defect and a value close to −1 to an under-twist defect (see an example in Fig. 1a).

In this section, we present the simulation results of both Snf2-naked DNA and Snf2-nucleosome systems, but focusing on the motion of the remodeler relative to the DNA. In Fig. 2a, we show that by switching between the remodeler chemical states during the MD simulation, this can slide along both naked and nucleosomal DNA by 1-bp (as evidenced by the change in the average lobe contact index during the ATP cycle). While sliding on naked DNA is not a key function of remodelers, this process has been documented in experiments(Sirinakis et al., 2011). Interestingly, we note that under the current computational settings sliding by 1 bp by the end of an ATP cycle occurs with higher probability when in complex with nucleosomes (98%) than when on naked DNA (45%), whereas in the remaining cases the remodeler simply goes back to its original position.

**Figure 2:**
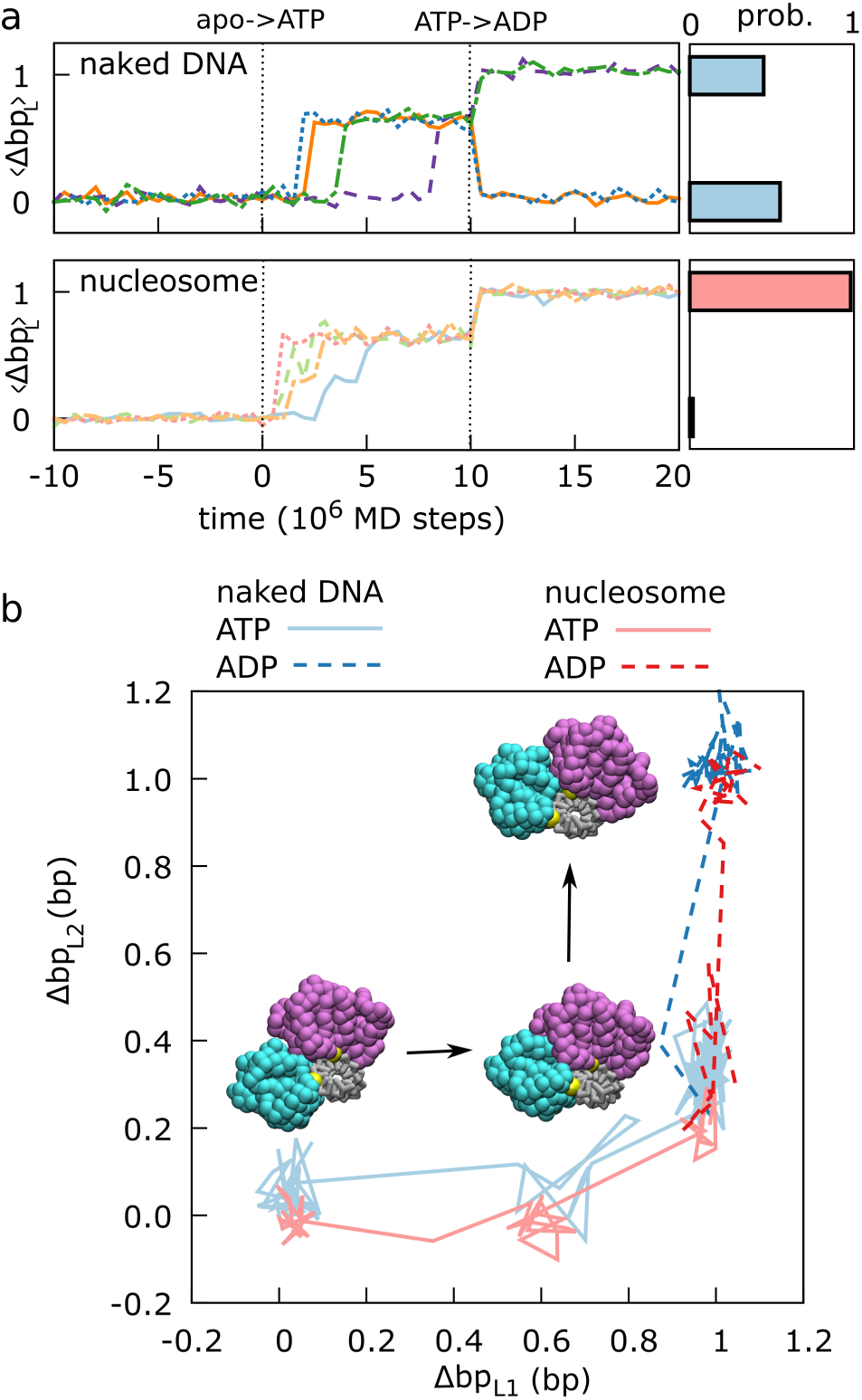
Snf2 translocase motion relative to DNA via an inchworm mechanism. (a) Representative trajectories of the average lobe contact index <Δbp_L_>=(Δbp_L1_+Δbp_L2_)/2 during the ATP cycle in our MD simulations (ATP binding occurs after equilibration at time 0, hydrolysis occurs after 10^7^ MD steps) on naked (upper left) and nucleosomal (lower left) DNAs with polyApG sequence. In both cases we find unidirectional motions; a 1 bp step of the translocase in direction from lobe 1 to lobe 2 occurs every ATP cycle with probabilities of ~0.45 and ~0.98 on naked DNA (upper right) and on nucleosomal DNA (lower right), respectively. (b) Projections of two representative trajectories (blue for naked, red for nucleosomal DNAs) on the lobe contact indexes Δbp_L1_ and Δbp_L2_ (lighter solid lines for the ATP state, darker dashed lines for the ADP state after hydrolysis), highlighting the inchworm mechanism. Snapshots of translocase moving on naked DNA are also shown (lobe 1 in cyan, lobe 2 in purple, DNA in gray, two reference phosphates in yellow).

Figure 2b displays two representative trajectories (one on the naked DNA and one on the nucleosome) projected onto the contact indexes of the two separate ATPase lobes. In both cases, this projection clearly highlights the inchworm motions. Specifically, starting from the apo state in the open conformation (bottom left in the figure), switching the potential to the ATP state induces the closure of the remodeler, with lobe 1 moving by 1 bp towards lobe 2, which maintains its position due to its stronger grip to the DNA (bottom right). Then, simulating ATP hydrolysis via switching to the ADP-state potential induces the domain opening, but since the lobe 2-DNA interactions are also decreased, now it is this lobe that usually moves by 1 bp away from lobe 1 (top right). Switching again to the apo-state potential simply restores the original lobes-DNA interaction strengths, maintaining the same open configuration and completing a full ATP cycle with the remodeler shifted by 1 bp relative to where it started. On naked DNA, this mechanism is sufficient to explain the translocase’s unidirectional motion (see Supplementary Movie 1 for a visualization of the trajectory in Fig. 2b). However, what is not clear from this analysis is how the translocase motion may induce sliding of nucleosomal DNA; the next sections are devoted to the characterization of the complete active repositioning process.

### Remodelers couple inchworm motion to nucleosome sliding via steric and electrostatic interactions

Our MD simulations show that the ATP-driven translocase closure is followed by sliding of nucleosomal DNA. Specifically, the DNA at the remodeler binding location slides unidirectionally towards the dyad, as indicated by the 1-bp increase in the average nucleosome contact index around SHL 2, (Δbp_1.5_+Δbp_2.5_)/2 (see two representative trajectories and the cumulative distribution at the end of the ATP-state in Fig. 3a, upper panel). This direction of repositioning is consistent with the experimental evidence(Liu et al., 2017). On the other hand, in the absence Snf2, sliding of the same polyApG nucleosomal DNA occurs in a random direction (Fig. 3a, lower panel).

**Figure 3:**
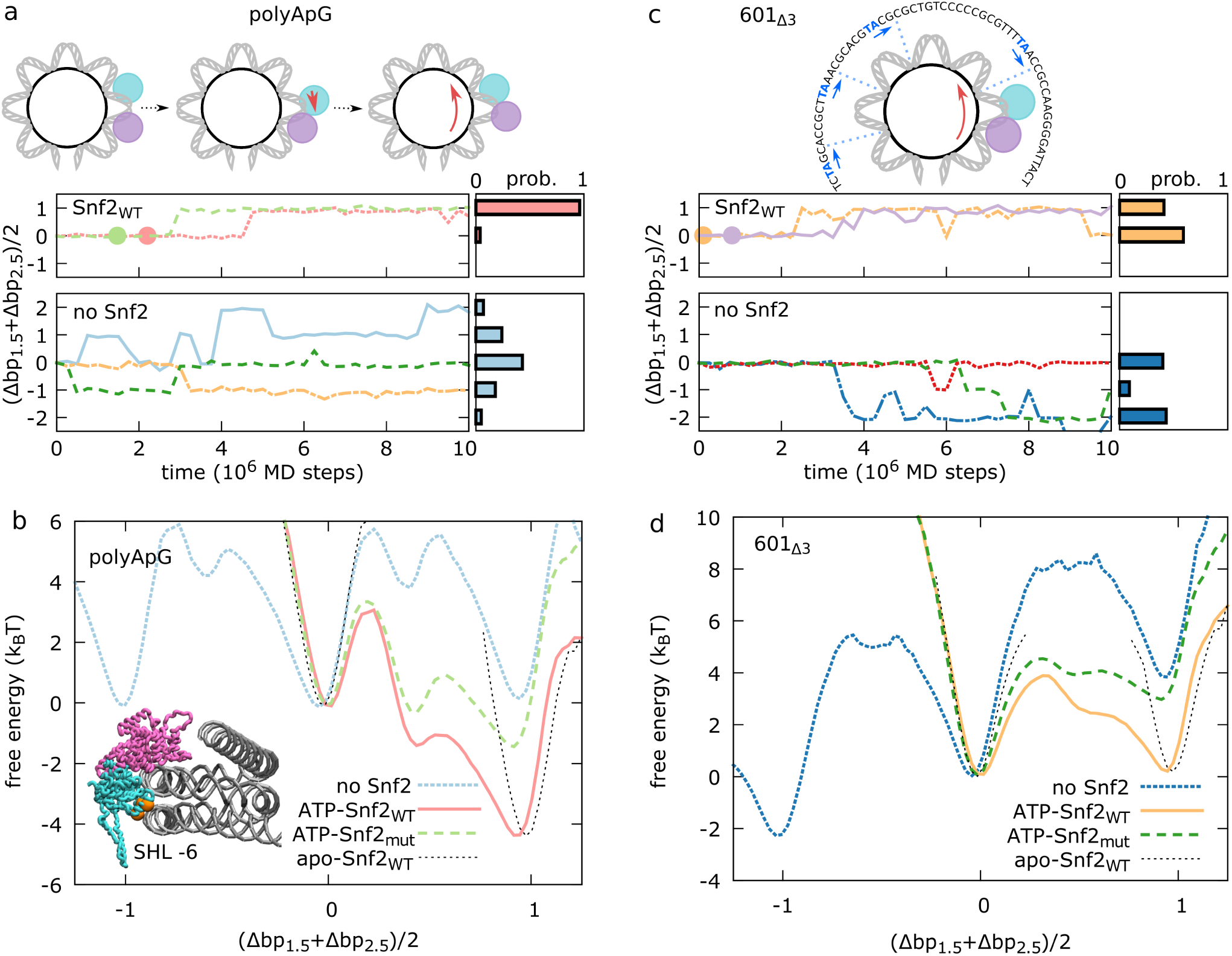
Mechanism of directed nucleosomal DNA sliding by Snf2. (a,c) For polyApG (a) and 601_Δ3_ (c) sequences, we show representative trajectories of the average contact index at SHL 2, (Δbp_1.5_+Δbp_2.5_)/2, as a function of time (left panel) and the repositioning histogram at the end of 100 trajectories of 10^7^ MD steps (right) for the remodeler-bound case (upper), and in the absence of remodeler (spontanous sliding) (lower). For the remodeler case, we report only the portion of trajectory in the ATP state, where nucleosome sliding occurs after the translocase closure (indicated by the circles). (b,d) For polyApG (b) and 601_Δ3_ (d) sequences we report the free energy profiles along the average contact index at SHL 2 for different systems: spontaneous sliding in the absence of the remodeler (no Snf2), in the complexes with the bound open apo-state Snf2 (apo- Snf2_WT_), the closed ATP-bound Snf2 (ATP-Snf2_WT_), and the closed ATP-bound K855E-R880E-K885E charge mutant Snf2 (ATP-Snf2_mut_). Errors on the free-energy profiles are on the order of ~0.3 k_B_T. The cartoons in panel a indicate how the inchworm motion of the translocase (lobe 1 in cyan, lobe 2 in purple) is coupled to nucleosome sliding. The structure of the Snf2-nucleosome complex in the inset in panel b highlights the location of the K855, R880 and K885 residues (orange) targeted by the mutation. In panel c we also displayed the central 63 base pairs of the 601_Δ_3 sequence highlighting the location of the TpA positioning steps within the Snf2-nucleosome complex (in blue). In the initial configuration for the 601_Δ3_ simulations, the DNA is shifted by 3 bp relative to the optimal structure with PDB id 5X0Y (with the TpA steps located where the DNA minor groove faces the histone octamer, indicated with a blue dashed line).

To better characterize the origin of the unidirectional motion, in Fig. 3b, we compare the free-energy profiles of nucleosome sliding along the average contact index around SHL 2 for different scenarios. In the absence of Snf2, as expected for the uniform polyApG sequence and the random motion reported in Fig. 3a, sliding by 1 base pair in either direction does not change the free energy of the system, but it involves climbing significant free-energy barriers (~6 k_B_T). The presence of the remodeler modifies the original nucleosome landscape in a chemical-state-dependent fashion. In the initial open conformation before ATP binding, there is a single free-energy minimum at (Δbp_1.5_+Δbp_2..5_)/2=0, so that nucleosomal DNA sliding is strongly inhibited. Instead, after ATP binding and translocase closure, a second deeper free-energy minimum appears around (Δbp_1.5_+Δbp_2.5_)/2=1, favoring DNA sliding towards the dyad. After the last opening conformational change following ATP hydrolysis, the nucleosome landscape returns to have a single free-energy minimum now at (Δbp_1.5_+Δbp_2.5_)/2=1, so that further sliding is inhibited. The switching among these different free-energy landscapes reveals a clear ratchet mechanism, as often employed for the theoretical modeling of molecular motors(Jülicher, Ajdari, & Prost, 1997). The changes in the free energy profiles can be in part explained by the inchworm motion of the translocase domain and steric effects. In the open conformation, DNA sliding at SHL 2 by 1 bp in either direction would cause steric overlap between lobe 2 and histone octamer on one side or overlap between lobe 1 and the opposite DNA gyre around SHL −6 on the other side (see inset in Fig. 3b), blocking nucleosome repositioning and explaining the single free energy minimum when Snf2 is in the apo state. Since the translocase closure upon ATP binding involves the motion of lobe 1 towards lobe 2, DNA sliding is now allowed to occur towards the dyad, causing the translocase to swing on the opposite side of its binding location (see cartoons in Fig. 3a). However, this argument does not explain the large extent to which the closed ATPase favors unidirectional repositioning, i.e. the decrease in free energy by ~4 k_B_T from (Δbp_1.5_+Δbp_2.5_)/2=0 to 1.

From the crystal structure of the nucleosome-bound Snf2 remodeler(Liu et al., 2017), it was shown that apart from the main interactions at SHL 2, the translocase domain also interacts with the opposite DNA gyre around SHL −6 via long-range electrostatics mediated by residues K855, R880 and K885, located within lobe 1 (see bottom view in Fig. 1c and inset in Fig. 3b). It was also experimentally shown that changing these residues from positively- to negatively-charged markedly reduced the remodeling activity of Snf2(Liu et al., 2017). To investigate this effect, we performed MD simulations where the three key residues have all been mutated to glutamic acid (K855E-R880E-K885E mutant). While still possible, DNA sliding around SHL 2 is no longer accompanied by a large decrease in free energy (Fig. 3b). This change can be understood in terms of the movement of the ATPase lobe 1 during repositioning. In the open state, lobe 1 is close to the contact point 1.5 and also interacts with the opposite gyre at SHL −6 via the basic patch in wild-type (WT) Snf2 (K855, R880 and K885). After ATP binding, lobe 1 moves by 1 bp towards lobe 2, becoming further apart from both contact point 1.5 and the DNA at SHL −6, weakening the electrostatic interaction (specifically, the average distance between the center of mass of the lobe 1 patch and the DNA phosphate backbone increases from ~6.1 Å to ~8.3 Å upon translocase closure). The sliding of nucleosomal DNA causes lobe 1 to swing back towards the dyad, restoring also the original interactions between the basic patch and SHL −6. Comparing to the initial open apo structure, the translocase closure and subsequent sliding of DNA at SHL 2 makes it appear that lobe 2 moved by 1 bp towards lobe 1, and not the opposite. Notably, this observation is consistent with the recent cryo-EM structure of the nucleosome-Chd1 complex in the presence of an ATP analog(Farnung et al., 2017), where lobe 1 overlaps with the corresponding lobe in the open conformation of the Snf2 remodeler(Liu et al., 2017), whereas lobe 2 appears to have moved by 1 bp(Farnung et al., 2017).

While so far we only considered a simple uniform polyApG sequence, genomes are rich in positioning motifs that contribute to specify the optimal location of nucleosomes along DNA(Struhl & Segal, 2013). These motifs, such as T/A base steps periodically spaced every 10 bp, cause the intrinsic bending of DNA, which lowers the free energy cost of nucleosome assembly, and favor a specific rotational setting, as they preferentially locate where the DNA minor groove faces the histone octamer(Freeman, Lequieu, et al., 2014). These signals strongly inhibit nucleosome sliding relative to random DNA sequences, since repositioning would proceed either by DNA screw-like motion via a high-energy intermediate with a non-optimal rotational setting(Brandani et al., 2018; Kulić & Schiessel, 2003), or via alternative repositioning mechanisms uncoupled with DNA rotations, which involve the energetically-costly breakage of many histone-DNA contacts(Lequieu et al., 2017; Niina et al., 2017). Nevertheless, chromatin remodelers are still able to actively reposition nucleosomes made with strong positioning sequences such as 601(Harada et al., 2016; Liu et al., 2017; Lowary & Widom, 1998).

To investigate the robustness of the active repositioning mechanism against changes in DNA sequence, we next run MD simulations of Snf2 in complex with nucleosomes made with the 601 sequence(Lowary & Widom, 1998). In the starting configuration, we shifted the DNA by 3 bp relative to the optimal configuration found in the 5X0Y structure, in the direction from the remodeler site towards the dyad. We refer to this sequence as 601_Δ3_. Because of the non optimal location of the T/A steps relative to the histone octamer (see cartoon in Fig. 3c), starting from here in the absence of the remodeler will be most likely followed by sliding backward away from the dyad, i.e. towards the optimal configuration (in about half of the cases within 10^7^ MD steps, Fig. 3c, lower panel). Instead, not only the remodeler prevents backward sliding, but upon ATP binding, in about half of the cases, it can also induce sliding of nucleosomal DNA forward towards the dyad (Fig. 3C, upper panel), in the same way as observed with the uniform polyApG sequence. A comparison of the free-energy landscapes along DNA sliding at SHL 2 with and without remodeler (Fig. 3d) shows indeed that in the case without remodeler the free energy strongly increases with sliding forward towards the dyad and decreases away from the dyad, whereas the closed translocase is able to lower the free energy cost of forward sliding to ~0 k_B_T, while preventing sliding backward in the opposite direction via steric effects. The free energy profile obtained with the K855E-R880E-K885E Snf2 bound to 601_Δ3_ nucleosomes, shows that this mutant cannot slide these strong positioning sequences, due to an extra free energy penalty of ~3 k_B_T upon sliding by 1 bp. This is consistent with the results from experiments on similar Snf2 charge mutants sliding 601 nucleosomes(Liu et al., 2017). While the limitations of our computational model (e.g. the assumptions on the precise ATP hydrolysis kinetics) prevent us from making quantitative predictions of remodeling activity, our simulations provide a mechanistic understanding of the important role of electrostatic interactions in directing repositioning(Liu et al., 2017).

### Nucleosome repositioning proceeds via the formation and propagation of twist defects from the remodeler binding location

So far, we focused on the inchworm motion of the translocase domain and on nucleosomal DNA sliding at the SHL 2 binding site, establishing how these two are tightly coupled. However, a full characterization of the repositioning mechanism requires the analysis of DNA sliding at the individual histone-DNA contact points on the entire nucleosome. In Fig. 4a, we plot the timelines of the contact index coordinates of both remodeler and nucleosome for two representative trajectories during which repositioning by 1 bp occurs. These plots show how nucleosomal DNA sliding is initiated near the translocase binding location at the contact point at SHL 1.5, with the creation of opposite-type twist defects at the neighboring SHLs. The diffusion of these defects then completes repositioning of the entire nucleosome. To aid the understanding of the dynamics, we label the key metastable conformations of the system according to the following rules: the first letter, o or c, corresponds respectively to open or closed translocase conformation; when it is closed (c), the domain can adopt distinct configurations with DNA and histone octamer within its binding site at SHL 2, which will be indicated by a capital letter as A, B, C or D (see below for definition); finally, a last integer number, 0, 1 or 2, indicates the number of over-twist defects which may form near the dyad at the three central SHLs (SHL −1, 0, and +1, these defects are most favorably found at SHLs +/-1).

**Figure 4:**
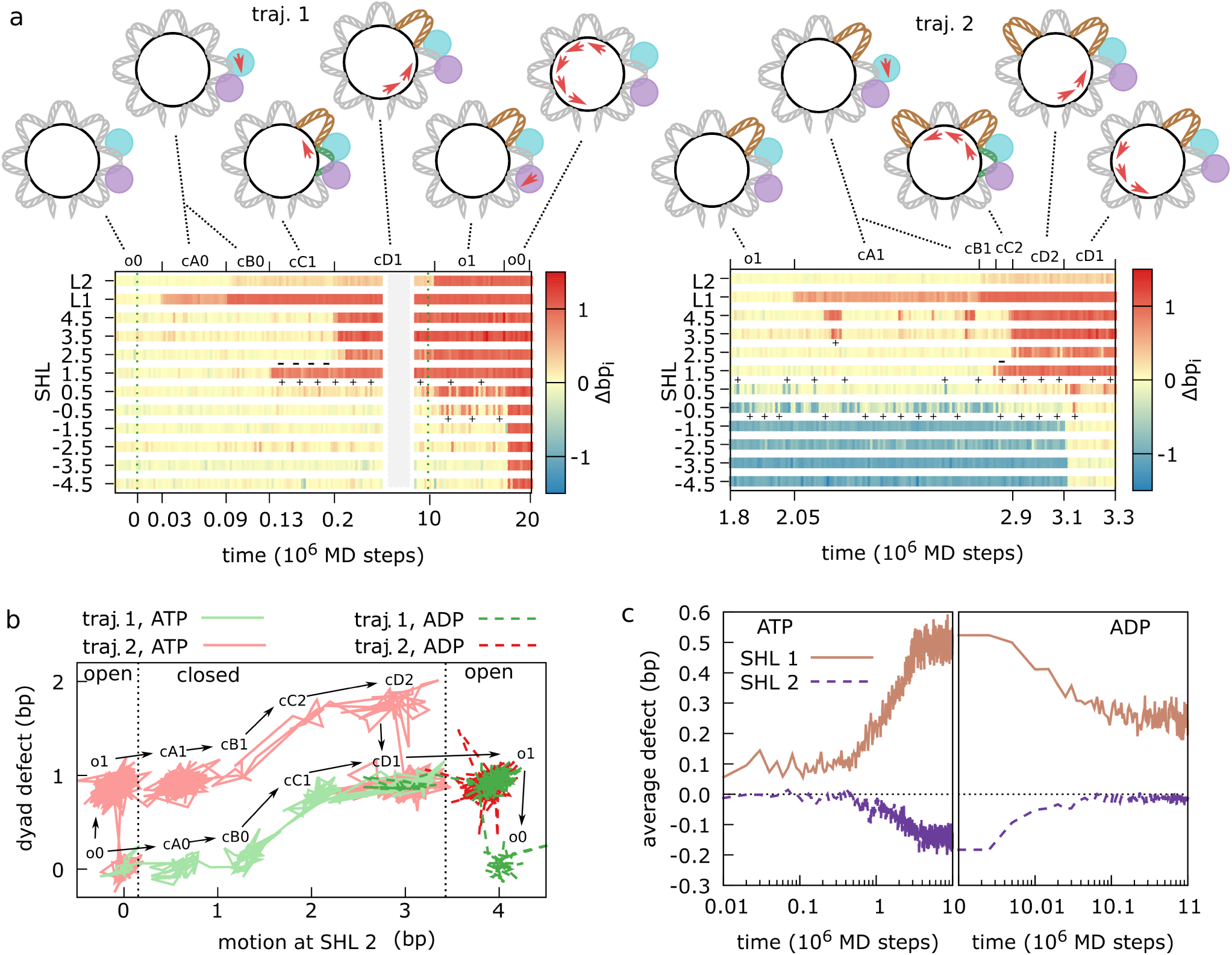
Active nucleosome repositioning via twist defect propagation for polyApG sequence. (a) Timelines of the translocase (L1 and L2) and nucleosome contact indexes (SHL −4.5 to 4.5) for two representative trajectories where nucleosomal DNA slides by 1 bp relative to the initial configuration. The intermediate configurations along the repositioning pathway are indicated by the corresponding labels (described in the main text) and cartoons. The key twist defects facilitating repositioning are highlighted by a plus sign for DNA over-twisting (in brown in the cartoons) and by a minus sign for DNA under-twisting (in green in the cartoons). DNA and translocase lobes (1 in cyan, 2 in purple) motions are indicated by red arrows. (b) 2-dimensional projections of the trajectories in panel a (traj. 1 in green, traj. 2 in red; lighter solid lines for the ATP state, darker dashed lines after hydrolysis; time increases in the direction indicated by the arrows). The x-axis represents the sum of the translocase and nucleosome contact indexes around the remodeler binding location at SHL 2: Δbp_L1_+Δbp_L2_+Δbp_1.5_+Δbp_2.5_. The y-axis represents the size of the twist defects at the three central SHLs: Δbp_1.5_-Δbp_-1.5_. (c) Twist defects at SHLs 1 (brown, solid line) and 2 (purple, dashed) averaged over 100 MD trajectories as a function of time, showing how twist defects are formed after ATP binding and translocase closure (left panel), and how they are released after hydrolysis and translocase opening (right panel).

In the first trajectory (the left panel in Fig. 4a), starting from an open translocase bound to a nucleosome in a standard 1KX5-like configuration lacking twist defects (state o0), switching to the ATP-bound potential at time 0 quickly induces the closure of the remodeler via the motion of lobe 1 towards lobe 2 (0.03×10^6^ MD steps, state cA0). In this first closed configuration, the lobe 1-DNA interface is destabilized relative to the one observed in the reference 5X0Y structure and the motion towards lobe 2 is only partial (Δbp_L1_~0.6). Only after some time (0.09×10^6^ MD steps) the motion of lobe 1 is complete (Δbp_L1_~1, state cB0). From this closed configuration, we observe motion of nucleosomal DNA towards the dyad relative to the histone octamer starting from SHL 1.5 (0.13×10^6^ MD steps, state cC1). In this configuration, the DNA motion causes DNA over-twisting at SHL 1, which now accommodates an extra base pair, and DNA under-twisting at SHL 2 (where the remodeler is bound), which now accommodates a missing base pair. Soon afterwards (0.2×10^6^ MD steps), the nucleosomal DNA further slides by 1 bp from the remodeler site up to the closest nucleosome entry/exit, releasing the under-twist defect (state cD1). As highlighted in the previous section, while these two steps do not involve remodeler’s motion relative to the DNA, the DNA motion relative to the histone octamer causes the remodeler to swing by 1 bp towards the dyad and enables to re-establish the electrostatic contacts between lobe 1 and SHL −6, which were lost during the initial ATPase closure. State cD1, for the polyApG sequence considered here, is the most stable configuration among the closed ones. Repositioning is usually completed only after ATP hydrolysis (10×10^6^ MD steps), which causes translocase opening via lobe 2 motion by 1 bp (state o1) (the full pathway is o0 → cA0 → cB0 → cC1 → cD1 → o1 → o0). The very last step consists of the sliding of nucleosomal DNA from the translocase up to the far nucleosome entry/exit, releasing the over-twist defect near the dyad (state o0, see Supplementary Movies 2 and 3 for visualizations of this trajectory). The second trajectory (the right panel) is qualitatively similar to the first, except that all states have an additional defect near the dyad at the starting time (the full pathway is then o1→ cA1→ cB1→ cC2→ cD2→ cD1→ o1 for trajectory 2). In particular, motion at the remodeler and nucleosome contact points proceeds in the same order. These two trajectories are representative of the most common repositioning pathways observed in our 100 MD trajectories. In all cases, repositioning involves twist-defect formation and propagation starting from the remodeler binding location.

Trajectories can be projected onto a low dimensional space defined by the sum of the contact indexes around the remodelers binding location (Δbp_L1_+Δbp_L2_+Δbp_1.5_+Δbp_2..5_, the horizontal axis in Fig. 3b) and by the size of the twist defect around the dyad (Δbp_1.5_-Δbp_-15_, the vertical axis in Fig. 4b). On this space, all the key metastable states involved in repositioning can be clearly separated (in Fig. 4b we show trajectories 1 and 2 from panel a). From this figure we notice that most key conformational changes occur in the closed conformation (between two dotted lines). To test the importance of the system relaxation in this portion of the phase space, we induced ATP hydrolysis after only 10^6^ MD steps from ATP binding (instead of 10^7^). Even if limiting our analysis to the trajectories where the remodeler successfully closed before hydrolysis (64%), only in 36% of the cases the remodeler can complete sliding by 1 bp, compared to the original 98% success rate (see Supplementary Fig. S1 for timeline of the contact indexes during one of these unsuccessful repositioning events). Failing to reposition occurs when the DNA does not have enough time to slide at the remodeler binding location (reaching state cD1), which is essential to avoid the steric overlap between lobe 2 and histones upon opening.

In Fig. 4c, we plot the changes in the average size of the twist defects at the SHLs near the remodeler during the ATP cycle. ATP binding and remodeler closure cause an enhancement of DNA over-twisting at SHL 1 and DNA under-twisting at SHL 2. Conversely, ATP hydrolysis and remodeler opening restore the former level of DNA twisting. Therefore, the closed remodeler lowers the free energy of opposite-type twist defects near its binding location at SHLs 1 and 2, which in turn will favor the initiation of DNA sliding towards the dyad starting from contact point 1.5, as observed in the trajectories in Fig. 4a.

### Defect-mediated repositioning is controlled by DNA sequence

The twist-defect-mediated mechanism highlighted by our MD simulations may offer a further route to control the remodeling activity via DNA sequence (apart from the effects of positioning motifs described above), potentially explaining the significant sequence-dependence observed in experiments(Brandani et al., 2018; Krietenstein et al., 2016; Lequieu et al., 2017; Lorch et al., 2014; Niina et al., 2017; Winger & Bowman, 2017). DNA sequence was already found to have a strong effect on the time-scales of spontaneous repositioning due to variations in DNA flexibility and energetics of twist defects(Brandani et al., 2018). When nucleosome diffusion is mediated by the formation of twist defects, repositioning proceeds much faster on sequences such as polyTpA, which are very flexible and easily accommodate twist defects, than on sequences such as poly(dA:dT) tracts (ApA repeats), which are stiffer and unlikely to display twist defects(Brandani et al., 2018). Perhaps surprisingly, it was shown that TpA repeats inhibit nucleosome repositioning by the Chd1 remodeler when the element is located at SHL 2(Winger & Bowman, 2017). The origin of the influence of sequence on active repositioning is not clear, and there is likely also a strong dependence on the considered remodeler itself, due to the many different domains that can interact with the nucleosome apart from the translocase(Clapier et al., 2017). We begin exploring the defect-mediated sequence effects on the behavior of our minimal remodeler system by considering targeted changes from reference polyApG sequence at specific regions: the insertion of a 10-bp poly(dA:dT) tract at SHL 1 (polyApG-ApA_SHL1_ sequence), and a 10-bp TpA repeat at SHL 2 (polyApG-TpA_SHL2_ sequence) (see Fig. 5a for the locations of these sequence elements within the Snf2-nucleosome complex). To characterize these effects, we reconstructed the free energy landscapes and kinetics of the systems via Markov state modeling(Prinz et al., 2011) (the details on this analysis are given in the Supplementary Information). In particular, we study the region of phase space where the translocase is in its closed (ATP-bound) configuration and lobe 1 fully completed the motion by 1 bp towards lobe 2, so that we can solely focus on the key formation and propagation of twist defects mediating repositioning.

**Figure 5:**
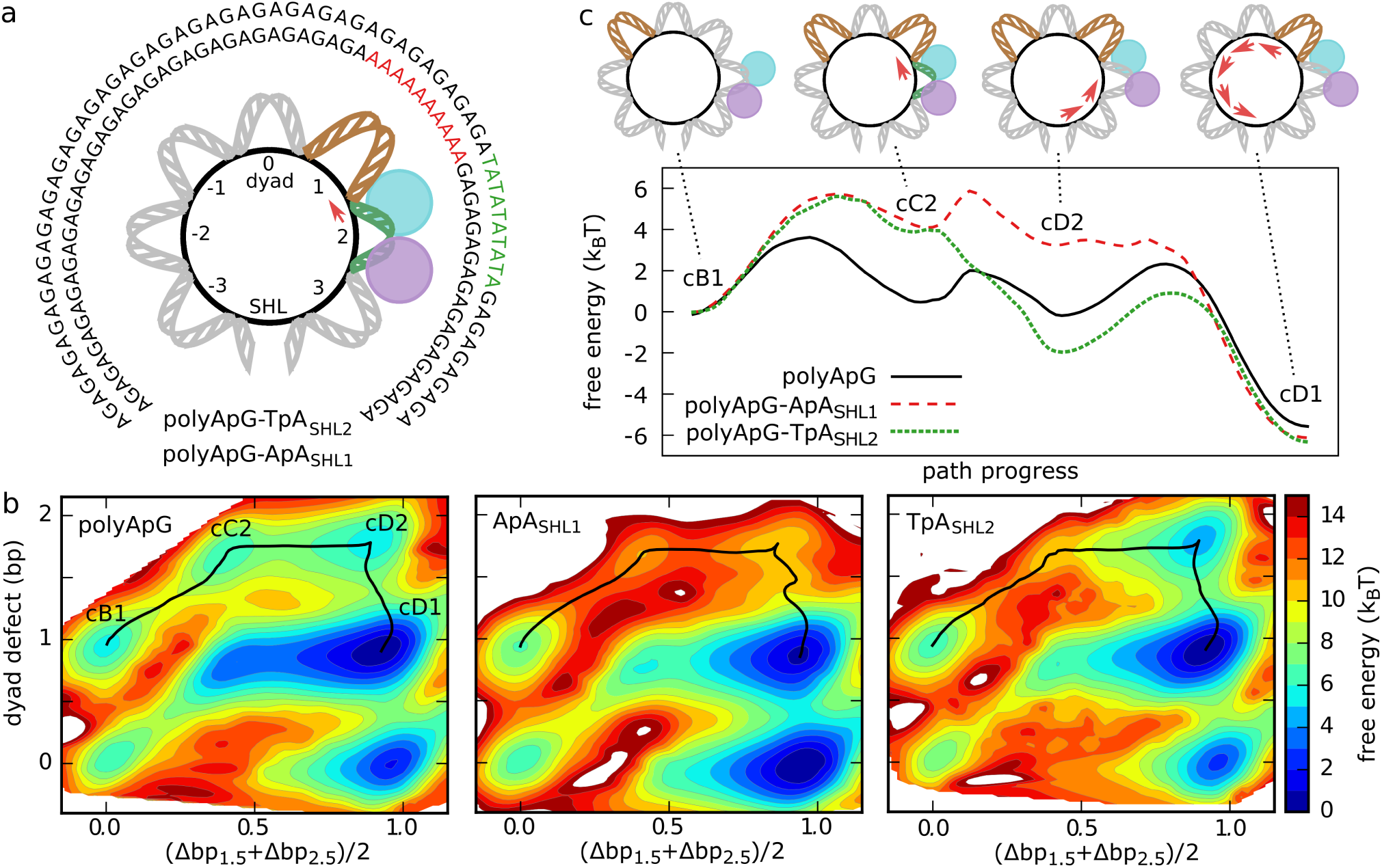
Twist-defect-mediated effects due to DNA sequence. (a) Considered DNA sequences apart from the reference polyApG: polyApG-ApA_SHL1_ (inner circle) and polyApG-TpA_SHL2_ (outer). We displayed the central 71 base pairs highlighting their location relative to the translocase-nucleosome complex. (b) Free energy landscapes of the Snf2-nucleosome complex in the closed conformation along the average contact index at SHL 2, (Δbp_1.5_+Δbp_2.5_)/2, and the size of the twist defects around the dyad (Δbp_1.5_-Δbp_-1.5_), for the three DNA sequences polyApG (left), polyApG-ApA_SHL1_ (center) and polyApG-TpA_SHL2_ (right). (c) Comparison of the free energy profiles along the minimum paths of repositioning from state cB1 to state cD1; polyApG (black, solid line), polyApG-ApA_SHL1_ (red, dashed), and polyApG-TpA_SHL2_ (green, dotted). The pathways are indicated on the landscapes in panel b by black solid lines. Errors on the profiles are within ~0.3 k_B_T.

In Fig. 5b, we display the free energy landscapes along the average contact index around SHL 2 (horizontal axis, (Δbp_1.5_+Δbp_2.5_)/2, the same coordinate used to investigate sliding in Fig. 3) and the number of twist defects at the three SHLs around the dyad (the vertical axis, Δbp_1.5_-Δbp**_-1.5_**, which enables to track how repositioning proceeds further away from SHL 2) for the three DNA sequences. These landscapes are qualitatively similar and have several local minima corresponding to the twist-defect intermediates observed in our trajectories. Indeed, the representative trajectories reported in Supplementary Fig. S2 shows how the role of twist defects in mediating active repositioning is robust against changes in DNA sequence and remodeler mutations. However, the changes considered here do have significant effects on the kinetics of repositioning. For instance, if we focus on the pathway of twist-defect propagation observed in trajectory 2 in Fig. 4 (cB1→cC2→cD2→cD1, which is the most common for polyApG, highlighted in black in Fig. 5b), then the addition of the ApA_SHL1_ and TpA_SHL2_ elements greatly increase the free energies of the intermediate states along the minimum energy pathways (Fig. 5c). This change is also evidenced by an increase in the mean first passage time to reach the final cD1 state (by a factor of ~2.4 and ~1.3 respectively). These changes are explained by the different DNA flexibilities and by the changes in defect pattern along the pathway of repositioning. ApA repeats where previously found to inhibit DNA over-twisting around the dyad, whereas TpA repeats where found to inhibit DNA under-twisting at SHLs +/-2(Brandani et al., 2018). Since states cB1 and cD1 have only one defect around the dyad (at SHL +1 or −1), whereas the intermediate states cC2 and cD2 have two defects (at both SHLs +1 and −1), the ApA_SHL1_ element increases the free energy of these two intermediate states. Similarly, despite its flexibility, the TpA_SHL2_ element significantly increases the free energy of the cC2 state, where DNA under-twisting at SHL 2 is observed. Therefore, the sequence-dependent formation and propagation of twist defects during remodeling provides an explanation of past experimental observations(Winger & Bowman, 2017).

Interestingly, we note that the repositioning activity may also depend on the DNA sequence at the translocase SHL 2 binding site via changes in the relative strength of lobe 1 and lobe 2 interactions with DNA. In particular, while the initial translocase closure occurring upon ATP binding always involves the motion of lobe 1 towards lobe 2 for polyApG, polyApG-ApA_SHL1_ and 601_Δ3_, in ~28% of the polyApG-TpA_SHL2_ simulations lobe 2 moves towards lobe 1, due to the weaker lobe 2-DNA interactions. When this occurs, there is no sliding of nucleosomal DNA, since the original lobe 1 electrostatic interactions with SHL −6 remain unaffected, and the translocase opening following ATP hydrolysis simply brings the system back to its starting configuration. This sequence effect resulting in lower remodeling activity on TpA repeats at SHL 2 is also consistent with past experiments(Winger & Bowman, 2017).

## DISCUSSION

The recent structural information of the Snf2-nucleosome complex(Liu et al., 2017) and insights from related molecular motors(Gu & Rice, 2010; Velankar et al., 1999) allowed us to design an efficient computational model that reproduces the expected inchworm motion of the translocase domain along DNA(Clapier et al., 2017). More importantly, our MD simulations revealed the detailed molecular mechanism by which the inchworm motion of the translocase is converted into nucleosome repositioning. Specifically, we establish the fundamental role of electrostatic interactions and nucleosome twist defects. The interaction between a basic patch located on lobe 1 and the DNA gyre around SHL −6 acts as an electrostatic spring to direct the sliding of nucleosomal DNA from the translocase binding site at SHL 2 towards the dyad. Firstly, ATP binding drives the closure of the translocase via the motion of lobe 1 towards lobe 2 by 1 bp. This initial motion increases the distance between lobe 1 and SHL −6, weakening the electrostatic interactions. The sliding of nucleosomal DNA towards the dyad then causes the translocase to swing towards dyad together with DNA, re-establishing the original electrostatic contacts. The steric repulsion between the remodeler and the histone octamer allows nucleosomal DNA sliding to occur in the closed conformation only, and prevents sliding in the opposite direction away from the dyad. This mechanism is robust enough to enable active repositioning of nucleosomes formed with strong positioning sequences such as 601(Lowary & Widom, 1998), where sliding has to proceed against a high free-energy uphill due to the preferred rotational register of T/A positioning motifs. Consistent with previous experiments, our MD simulations of Snf2 mutants show that the basic patch on lobe 1 is essential for the ability to slide 601 nucleosomes(Liu et al., 2017).

The electrostatics-driven sliding of nucleosomal DNA starts from the histone-DNA contact point at SHL 1.5, via the formation of a pair of twist defects of opposite type at the neighboring SHLs: DNA under-twisting at SHL 2 accommodates the missing base pair, DNA over-twisting at SHL 1 accommodates the extra base pair. The importance of twist defects lies in facilitating the initiation of nucleosomal DNA sliding locally from the translocase binding site, without simultaneously breaking distant histone-DNA contacts. The initial paired twist-defect conformation represents an intermediate state with tension accumulated around the remodeler location. Release of this tension via the propagation of the two twist defects in opposite directions completes the sliding of DNA by 1 bp throughout the entire nucleosome. The ATP cycle is completed after hydrolysis and ADP release, during which the motion of the weakened lobe 2 away from lobe 1 brings the translocase in the initial open conformation, only with the nucleosomal DNA slid by 1 bp relative to both histone octamer and remodeler. Interestingly, the twist defects that favor active nucleosome repositioning are the same that are commonly found in nucleosome crystal structures (at SHL 2)(Luger et al., 1997; Richmond & Davey, 2003), and those that have been shown to play an important role in spontaneous repositioning (SHL 1 and 2) (Brandani et al., 2018), suggesting a reason for targeting the specific translocase binding location.

Our results are consistent with the wave-ratchet-wave model of nucleosome repositioning(Clapier et al., 2017; Saha et al., 2005), where sliding is initiated at the translocase site via the generation of tension (DNA twisting). This has been suggested as a unifying fundamental mechanism of chromatin remodeling, where additional domains control tension release and interactions with other nucleosome regions to perform substrate recognition and to define the remodeling outcome (e.g. sliding vs histone ejection vs histone exchange)(Clapier et al., 2017). While we considered a minimal remodeler system consisting of nucleosome and translocase domain only, the molecular mechanism highlighted here may offer an understanding of more complex scenarios. For instance, the ISWI remodeler was found to slide nucleosomes via coordinated 3-bp entry and 1- bp exit steps(Deindl et al., 2013). These findings are consistent with our model if the additional HAND-SANT-SLIDE domain of ISWI blocks DNA sliding on the entry side until enough tension from under-twist defects is accumulated (3 bp), while on the exit side DNA sliding is unrestrained and it proceeds at 1-bp steps via the release of over-twist defects(Clapier et al., 2017). Although our computational model is currently not well tested for the study of histone conformational changes, the strong accumulation of tension from DNA twisting on the entry side may be related to the histone octamer distortions that are necessary for the remodeling activity of ISWI(Sinha et al., 2017). Distortions at the same histone regions have also been shown to contribute to thermally-driven nucleosome sliding(Bilokapic, Strauss, & Halic, 2018), and their role deserves further investigation. Future studies should also address how remodelers may couple sliding of nucleosomal DNA to histone exchange(Brahma et al., 2017).

Finally, our simulations reveal how the twist-defect mechanism allows a kinetic control of nucleosome repositioning via DNA sequence. This is due to the significant sequence-dependence in the formation of the twist-defect intermediates that facilitate repositioning. In our previous work (Brandani et al., 2018), we showed how the free-energy cost of DNA over-twisting and under-twisting changes as a function of superhelical location (SHL) and sequence. With this knowledge, it is possible to target precise remodeling intermediates to control nucleosome sliding. Specifically, we tested the effect of introducing a 10-bp TpA repeat at SHL 2, which inhibits under-twist defects at this location, and a 10-bp poly(dA:dT) tract at SHL 1 (ApA_SHL1_), which inhibits over-twist defects. Consistent with the important role of twist defects at these regions, both DNA sequence modifications slow-down sliding relative to the pure polyApG case, due to the increase in the free-energy barrier along the repositioning pathway. Notably, our TpA simulations explain recent experiments showing that the addition of this sequence element blocks repositioning by the Chd1 remodeler at SHL 2(Winger & Bowman, 2017). These results may be considered surprising, because TpA base steps are known for their high flexibility(Olson et al., 1998) and expected to favor nucleosome sliding. Indeed, polyTpA sequences were found to be among those with the highest spontaneous nucleosome diffusivity(Brandani et al., 2018). However, even in that case, DNA sliding at SHL 2 represented the main kinetic bottleneck to sliding due to the high twist-defect cost(Brandani et al., 2018), further supporting the tight relationship between spontaneous and active scenarios. Many experiments have also highlighted the role of poly(dA:dT) tracts in controlling remodeling outcomes(Krietenstein et al., 2016; Lorch et al., 2014; Winger & Bowman, 2017). Of note, while these sequences are well known for inhibiting their wrapping into nucleosomes(Segal & Widom, 2009), recent experiments suggested that the increase in the free energy of nucleosome assembly is not sufficient to explain the formation of nucleosome free regions(Lorch et al., 2014), and that these are instead created via the direct action of chromatin remodelers(Krietenstein et al., 2016; Lorch et al., 2014). Our molecular dynamics simulations reveal the detailed molecular mechanism by which DNA sequence changes affecting twist-defect formation may be exploited to control remodeling activities.

We believe that the predictions of our simulations, such as the existence of distinct twist defect patters induced by the remodeler, could be potentially tested via Cryo-EM, which is becoming more and more accurate in the characterization of the structural heterogeneity of biomolecular complexes(Bilokapic et al., 2018; Fernandez-Leiro & Scheres, 2016). Finally, our computational methods may be readily employed for investigating the activity of different remodelers with known structural information, such as Chd1 (Farnung et al., 2017) and INO80(Ayala et al., 2018; Eustermann et al., 2018), and for studying remodeling in the biologically-relevant context of multi-nucleosome chromatin fiber(Nikitina, Norouzi, Grigoryev, & Zhurkin, 2017).

## MATERIALS AND METHODS

Our coarse-grained model employs the AICG2+ structure-based force field for proteins(Li et al., 2014) and the 3SPN.2C force field for DNA(Freeman, Hinckley, et al., 2014). This combination has been successfully applied in many past studies of nucleosome dynamics(Brandani et al., 2018; Chang & Takada, 2016; Freeman, Lequieu, et al., 2014; Kenzaki & Takada, 2015; Lequieu et al., 2017; Niina et al., 2017), and it enables a good compromise between computational speed-up and accuracy, necessary to reach the time-scales relevant to nucleosome repositioning(Brandani et al., 2018; Lequieu et al., 2017; Niina et al., 2017). According to these models, each amino acid is coarse-grained to a single bead located at the corresponding Cα atom(Li et al., 2014), whereas each nucleotide is represented by 3 beads corresponding to sugar, phosphate and base groups(Freeman, Hinckley, et al., 2014). The reference native conformations of the histone octamer and the Snf2 ATPase are respectively taken from the nucleosome crystal structure with PDB id 1KX5(Davey et al., 2002) and the nucleosome-Snf2 cryo-EM structure with PDB id 5X0Y(Liu et al., 2017). Histone tails and remodeler disordered regions not visible in the reference structures are modeled according to a sequence-dependent local statistical potential(Terakawa & Takada, 2011). The DNA model was parametrized against several experimental thermodynamic quantities, such as melting temperature,(Freeman, Hinckley, et al., 2014) and the sequence-dependent elasticity of DNA(Olson et al., 1998). Notably, this model has been successfully employed to predict the role of DNA sequence on the free energy of nucleosome assembly(Freeman, Lequieu, et al., 2014) and the formation of twist defects(Brandani et al., 2018), suggesting its suitability for investigating the sequence-dependent effects present in chromatin remodeling(Krietenstein et al., 2016; Lorch et al., 2014; Winger & Bowman, 2017).

Histone octamer, remodeler and DNA interact via excluded volume, long-range electrostatics and hydrogen-bonds. For excluded volume, we employ bead-type dependent radii derived from a database of protein-protein and protein-DNA complexes(Brandani et al., 2018; Niina et al., 2017; Tan, Terakawa, & Takada, 2016). Following our previous protocol(Brandani et al., 2018; Niina et al., 2017), electrostatics is modeled according to Debye-Hückel theory, with standard unit charges placed on DNA phosphate groups and protein residues in flexible regions, and fractional charges on protein residue in folded regions derived using the RESPAC method, which optimizes the coarse-grained electrostatic potential against the all-atom one in the folded conformation(Terakawa & Takada, 2014). For intra-DNA electrostatics, phosphate charges are rescaled by 0.6 to implicitly account for counter ion condensation(Freeman, Hinckley, et al., 2014). In all simulations the salt concentration was set to 250 mM of monovalent ions. Histone-DNA hydrogen bonds are modeled using a recently-developed distance- and orientation-dependent potential between protein residues and phosphate groups(Brandani et al., 2018; Niina et al., 2017); the potential, unlike a Go-like contacts(Kenzaki & Takada, 2015), is invariant under a rotation-coupled motion of DNA, allowing the study of nucleosome repositioning. These hydrogen bonds are defined from those observed in the 1KX5(Davey et al., 2002) and 3LZ0(Vasudevan, Chua, & Davey, 2010) crystal structures(Brandani et al., 2018; Niina et al., 2017), and we employed the same hydrogen bond strength ε_HB_ = 1.8 k_B_T used in Ref.(Brandani et al., 2018), which is an intermediate value among those giving a nucleosome disassembly profile consistent with experiments(Niina et al., 2017). Properly accounting for hydrogen-bond interactions is necessary for reproducing the experimental twist-defect metastability playing an important role in spontaneous nucleosome repositioning(Brandani et al., 2018; Kulić & Schiessel, 2003).

Translocase-DNA contacts are identified from the protein hydrogen donors within 5 Å from phosphate-group oxygen acceptors in the 5X0Y Snf2-nucleosome structure, and are represented with the same potential used for histone-DNA hydrogen bonds (reference distances and angles are also taken from 5X0Y). With this choice, lobe 1 (residue id 743-940) and lobe 2 (residue id 1046-1212) have each 16 contacts with DNA. These interactions are not strictly speaking only hydrogen bonds, but contacts introduced in the coarse-grained model to specify the remodeler-DNA binding mode. The relatively high cutoff is necessary to ensure that the motion of the remodeler lobes relative to the DNA occurs only during the translocase opening or closure steps induced during the ATP cycle, since spontaneous remodeler motion in the absence of ATP is inconsistent with the inchworm mechanism suggested from experiments(Clapier et al., 2017; Gu & Rice, 2010; Velankar et al., 1999). To model the remodeler in the closed conformation adopted upon ATP binding, we created additional lobe 1-lobe 2 Go-like contacts identified from the Cα atoms within 7.5 Å in a homology model generated after aligning the two Snf2 lobes to the corresponding lobes found in the closed conformation of the NS3 helicase with PDB id 3KQU(Gu & Rice, 2010). This reference structure was chosen because it was shown that the apostate open conformations of Snf2(Liu et al., 2017) (PDB id 5X0Y) and NS3(Gu & Rice, 2010) (PDB id 3KQH) display a high degree of structural similarity(Liu et al., 2017).

As described at the beginning of the Results section, we simulate the remodeling activity of Snf2 during a full ATP cycle by switching the underlying potential to model changes in the translocase chemical state, as previously done for modeling other protein motors(Koga & Takada, 2006; Yao et al., 2010). Each MD simulation cycle consists of 2×10^7^ MD equilibration steps in the apo-state, 10^7^ steps in the ATP-bound state, and 10^7^ steps in the ADP-bound state. In the apo state, translocase-DNA contacts have the same strength as histone-DNA hydrogen bonds (ε_1-__dna_ = ε_2-__dna_ = 1.8 k_B_T) and lobe 1-lobe 2 contacts are very weak (ε_1-2_ = 0.17 k_B_T), so that the other native interactions based on the 5X0Y reference dominate and the translocase is preferably in its open conformation. In the ATP-bound state, translocase-DNA interactions remain unchanged, but the interactions between lobe 1 and lobe 2 are strengthened (ε_1-2_ = 1.0 k_B_T) to stabilize the closed conformation. After ATP hydrolysis, lobe 2-DNA interactions are weakened by a factor of 0.8 (ε_2-DNA_ = 1.44 k_B_T). Completing the cycle requires a last switch to the apo-state potential, but this only changes the translocase- DNA interactions, without inducing any conformational change, therefore this step is not shown in the results. With these settings, we generate with very high probability successful nucleosome repositioning by 1 bp in the correct direction(Liu et al., 2017) (DNA sliding from the remodeler site towards the dyad, see Results section) via the inchworm mechanism suggested from past experimental studies(Clapier et al., 2017; Gu & Rice, 2010; Velankar et al., 1999). Variations to this simulation protocol, such as using the recent nucleosome-Chd1 cryo-EM structure to model the ATP-bound closed conformation(Farnung et al., 2017) or changes to the Snf2-DNA interactions, have been also tested, but they do not affect the key features of the active repositioning mechanism mediated by twist defects presented in the Results section.

MD simulations have been performed using the software CafeMol 3.0(Kenzaki et al., 2011) (available at http://www.cafemol.org/), integrating the equations of motion using the default settings via Langevin dynamics at 300 K. For the cases where the remodeler is in complex with the nucleosome, we ran 100 MD simulation cycles for each system, whereas for the remodeler sliding on naked DNA we ran 40 cycles. In the former cases, nucleosomal DNA is made by 223 bp (central 147 bp plus 38 bp for each linker), whereas in the latter by 40 bp. Nucleosome-bound simulations start from the conformation observed in the 5X0Y structure(Liu et al., 2017) after a short energy minimization with the steepest descent method (Fig. 1 c). In the Supplementary Information we provide details on the generation of the free-energy landscapes and kinetics of the systems from our MD trajectories via Markov state modeling(Prinz et al., 2011).

## ACKNOWLEDGEMENTS

We thank Ralf Blossey for stimulating discussion on chromatin remodeling mechanisms. We are also grateful to Tsuyoshi Terakawa for carefully reading our manuscript. This work was supported by JSPS KAKENHI grants [25251019 to S.T., 16KT0054 to S.T., and 16H01303 to S.T.], by the MEXT as “Priority Issue on Post-K computer” (S.T.), and by the RIKEN Pioneering Project “Dynamical Structural Biology” (S.T.). The funders had no role in study design, data collection and analysis, decision to publish, or preparation of the manuscript.

